# Analysis and Identification of Necroptosis Landscape on Therapy and Prognosis in Bladder Cancer

**DOI:** 10.1101/2022.05.11.491452

**Authors:** Zihan Zhao, Ning Jiang, Yulin Zhang, Yuhao Bai, Tianyao Liu, Tianhang Li, Hongqian Guo, Rong Yang

## Abstract

Bladder cancer (BLCA) is one of the most common malignant tumors of the urinary system, but current therapeutic strategy based on chemotherapy and immune checkpoint inhibitors (ICIs) therapy cannot meet the treatment needs, which mainly owing to the endogenous or acquired apoptotic resistance of cancer cells. Targeting necroptosis provides a novel strategy for chemotherapy, targeted drugs, and improves the efficacy of ICIs because of strong immunogenicity of necroptosis. Therefore, we systemically analyzed necroptosis landscape on therapy and prognosis in BLCA. We firstly divided BLCA patients from The Cancer Genome Atlas (TCGA) database into two necroptosis-related clusters (C1 and C2). Necroptosis C2 showed a significantly better prognosis than C1, and the differential genes of C2 and C1 were mainly related to the immune response according to GO and KEGG analysis. Next, we constructed a novel necroptosis related genes (NRGs) signature consisting of SIRT6, FASN, GNLY, FNDC4, SRC, ANXA1, AIM2, and IKBKB to predict the survival of TCGA-BLCA cohort, the accuracy of NRGsocre was also verified by external datasets. In addition, a nomogram combining NRGscore and several clinicopathological features was established to predict the BLCA patient’s OS more accurately and conveniently. We also found that NRGscore was significantly related to the infiltration levels of CD8 T cells, NK cells, and iDC cells, the gene expression of CTLA4, PD1, TIGIT, and LAG3 of TME, the sensitivity to chemotherapy and targeted agents in BLCA patients. In conclusion, the NRGscore has an excellent performance in evaluating the prognosis, clinicopathologic features, tumor microenvironment (TME) and therapeutic sensitivity of BLCA patients, which could be utilized as a guide for chemotherapy, ICIs therapy, and combination therapy.

## 1 Introduction

Bladder cancer (BLCA) is one of the most common malignant tumors of the urinary system, and its mortality and morbidity are increasing in recent years (1). According to the latest data from the National Cancer Center of China, BLCA ranks eighth in the number of new cancer cases among Chinese men, with an estimated number of 62,000 cases and 25,000 deaths (2). Although the majority of new cases are non-muscular invasive tumors, the 5-year survival rate is 90%. Once the muscular infiltration stage is entered, the 5-year survival rate drops to 36-48%, with only 5-36% in advanced patients (3). At present, the first-line treatment for advanced bladder cancer is dominated by combination chemotherapy with cisplatin, with an objective response rate of 50-60%. However, due to the high heterogeneity of BLCA, about 40% of patients are still unable to benefit from it or even suffer continuous progression (4). Due to the strong immunogenicity in tumor microenvironment (TME) of BLCA, the immune checkpoint inhibitors (ICIs) have also achieved certain therapeutic effects in the treatment of BLCA (5). Although the therapeutic effect of immune checkpoint inhibitors (ICIs) in BLCA better than other cancers, it still faces the problem of less than 30% response rate (6). It is necessary to optimize the currently used therapy strategy, including chemotherapy and ICIs, before the next generation of effective antitumor treatment regimens are mature, such as adoptive cell therapy.

Whether it is cisplatin-based chemotherapy or ICIs represented by anti-PD-1/PD-L1, the main way to effectively kill tumor cells is to induce tumor cell apoptosis. Previous study shown that even among these 50-60% of patients who benefit from chemotherapy, there are still a lot of patients will develop resistance to cisplatin during treatment, which may own to the resistance of tumor cells to apoptosis (7). In response to the resistance generated by ICIs, recent studies have shown that the immunogenicity of tumors can be improved through combination therapy, thereby increasing the sensitivity of tumors to immunotherapy (8). Therefore, it is a feasible strategy to improve the therapeutic effect of chemotherapy and immunotherapy in BLCA by developing methods to induce non-apoptotic forms of programmed cell death as a combination therapy.

Necroptosis is a novel apoptosis-independent programmed cell death with strong immunogenicity that plays a critical role in cancer patient prognosis, cancer progression and metastasis, cancer immune surveillance, and cancer subtypes important role (9). The lower expression of the key molecular in necroptosis signaling pathway was associated with the poor prognosis and promote the tumorigenesis and metastasis in multiple cancers (10-14). Besides, in clinical trials of immune checkpoint inhibitors therapy, the key molecules of necroptosis, including RIPK1, RIPK3 and MLKL, were associated with the function of T cells, and the overexpression of these genes was related to the better survival (15). The NF-κB signaling derived from necroptosis can enhance CD8+ T cell-mediated tumor killing by activating dendritic cells and increasing the tumor antigens presentation (16). In addition, considering the tolerance of tumor cells to apoptosis is responsible for the resistance to chemotherapy, induction of necroptosis acts as an effective strategy to address the issue of apoptosis resistance during chemotherapy and a variety of anticancer drugs have been developed to induce necroptosis (17). Totally, given the critical role of necroptosis in cancer development and treatment, necroptosis has emerged a great potential for cancer therapy by improving the therapeutic effect of chemotherapy and immunotherapy in BLCA.

However, there are few studies focusing on necroptosis of BLCA as far. In this study, we analyzed the expression of necroptosis-related genes (NRGs) in 335 BLCA patients from The Cancer Genome Atlas (TCGA) database. The survival analysis, TME analysis, and treatment efficacy analysis of NRGs were used. Moreover, a prognostic signature (NRGscore) and nomogram model based on NRGscore were constructed. The validation with independent cohort showed that the NRG score provided accurate prognostic prediction for BLCA patients. Considering, we have verified that NRGscore is significantly correlated with TME and prognosis of BLCA patients, which suggests that targeting necroptosis through various drugs that induce or manipulate necrotic pathways in BLCA may be considered as an adjuvant therapy strategy for BLCA patients.

## 2 Materials and Methods

### 2.1 Access to Datasets and Patients Information

**2.2** Transcription RNA sequencing and clinical information of patients in BLCA cohort were publicly obtained from the Cancer Genome Atlas (TCGA) database (https://portal.gdc.cancer.gov/). Transcription RNA sequencing included 414 BLCA tissues and 19 normal bladder tissues. The RNA-sequencing data in this cohort was downloaded as fragments per kilobase of transcript per million mapped reads (FPKM). When an individual gene symbol contained more than one Ensembl ID in RNA sequencing data, gene expression was annotated averagely. We removed 98 samples with incomplete clinical information, and finally 335 patients were included in subsequent study. We used independent cohorts (GSE13507 and GSE31684) containing merged 258 primary bladder cancer samples and for external validation. The corresponding RNA-seq data and the survival information of patients were obtained from the GEO (http://www.ncbi.nlm.nih.gov/geo).

### 2.3 Retrieval of Necroptosis-Related Genes

We obtained 598 necroptosis-related genes (Supplementary Table S1) from the genecard (https://www.genecards.org/). Using univariate Cox analysis, 17 genes associated with bladder cancer prognosis (p<0.01) were identified and used as NRGs for subsequent analysis.

### 2.4 Consensus clustering for NRGs and Functional Enrichment Analysis

Based on the gene expression of 17 NRGs in TCGA cohort, unsupervised clustering analysis was applied to determine the number of clusters. Finally, consensus clustering analysis in TCGA cohort identified two clusters using the “ConsensusClusterPlus” package. The “ggplot2” and “limma” package were used to construct a principal component analysis (PCA). We analyzed clinicopathological differences between the two clusters with the Chi-square test. The “survival” package was applied to compare survival differences between two clusters. Gene ontology (GO) and Kyoto Encyclopedia of Genes and Genomes (KEGG) analyses were performed by the “clusterProfiler”, “org.Hs.eg.db”, “enrichplot” and “GOplot” to conduct functional annotation of the genes with different expression (false discovery rate (FDR) < 0.05 and log2 fold change (FC) > 1) in two clusters. Analysis results are visualized through the “ggplot2” package.

### 2.5 Construction and Validation of NRGs Prognostic Signature

Based on 17 bladder prognosis-related NRGs that have been obtained by univariate Cox analysis, we performed the least absolute shrinkage and selection operator (LASSO) by “glmnet” package. Subsequently we used “survival” and “MASS” package in R to choose a model with multivariate Cox analysis which was operated in a stepwise algorithm filtered by AIC value to identify NRGs risk signature. The NRGs risk score formula was obtained as follows:

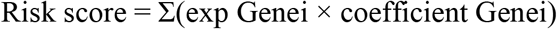

### 2.6 Patients in TCGA cohort were classified into high-risk and low-risk groups based on the median score

We used “survival” and “survminer” package to perform Kaplan Meier (K-M) analysis. Package “timeROC” in R was used for the analysis of receiver operating characteristic (ROC) curve and to calculate the area under the ROC curves (AUCs). We used GSE13507 and GSE31684 datasets as the validation cohort to validate the prognostic value of the NRGs risk signature. Patients risk scores were calculated using the same formula as above and the same cutoff criteria was used to classify the patients into low-risk and high-risk groups. Then, Kaplan–Meier survival analysis and ROC curve analysis were performed in validation cohort to assess the prognostic value. Clinicopathological data differences between the two risk groups were assessed by the Chi-square test and presented by a heatmap. Besides, univariate and multivariate Cox proportional hazards regression analyses were performed to evaluate whether the risk score and clinicopathological variables were independent prognostic factors for overall survival.

### 2.7 Establishment of the Nomogram Model

We use NRGs risk scores and clinicopathological factors to build a nomogram model to provide more accurate prognosis prediction for BLCA patients. The “rms” package in R helped to establish a nomogram for predicting prognosis. Also, we established calibration curves to test whether predicted survival was consistent with actual survival by “rms” package.

### 2.8 Gene Set Enrichment Analysis Between NRGs Risk Groups

Gene Set Enrichment Analysis (GSEA, http://www.broad.mit.edu/gsea/) is a computational method that determines whether an a priori defined set of genes shows statistically significant, concordant differences between two biological states. We performed GO-GSEA and KEGG-GSEA analyses between high-risk and low-risk groups using GSEA software (version 4.1.0). We chose gene set databases as “pub/gsea/gene_sets/c5.go.v7.5.1.symbols.gmt” and “pub/gsea/gene_sets/c2.cp.kegg.v7.5.1.symbols.gmt”.

### 2.9 Immune Checkpoint and Immune Enrichment Analysis

We explored differential expression of immune checkpoints between high-risk and low-risk groups, which may be related with the treatment responses of immune checkpoint inhibitors. In order to assess the tumor immune microenvironment (TIME) status of BLCA, we used single sample gene set enrichment analysis (ssGSEA) in R by “GSVA”, “Limma” and “GSEABase” package to evaluate a total of 16 congenital and adaptive immune cells as well as 13 immune-related functions.

### 2.10 Evaluation of the Efficacy of Treatment Response

We explored the value of the predictive signature in predicting the response to BLCA treatment. The “pRRophetic” package was used to perform the ridge regression algorithm to calculate the half-maximal inhibitory concentration (IC50). We also assessed differences in the response to immunotherapy between high-risk and low-risk groups. We uploaded the BLCA patients’ gene expression data to the TIDE website (http://tide.dfci.harvard.edu/) and obtained the TIDE score of each patient. We also obtained immunophenoscore of TCGA BLCA patients from the TICA website (https://www.tcia.at/home).

### 2.11 Statistical Analysis

R software (version 4.1.3) and GSEA software (version 4.1.0) were used for all statistical analysis and diagram drawing. Univariate Cox analysis was used to identify NRGs associated with overall survival (OS). Lasso regression analysis and multivariate Cox analysis filtered by AIC values were used to construct a predictive signature. The survival of patients in the high-risk and low-risk groups was analyzed by Kaplan-Meier method and log-rank test. Wilcoxon test was to identify the significance of the difference in two risk groups. P-value <0.05 was set as a statistically significant standard.

## 3 Results

To describe our research intuitively and systematically, we present the research process in **Figure 1**.

**Figure 1.**
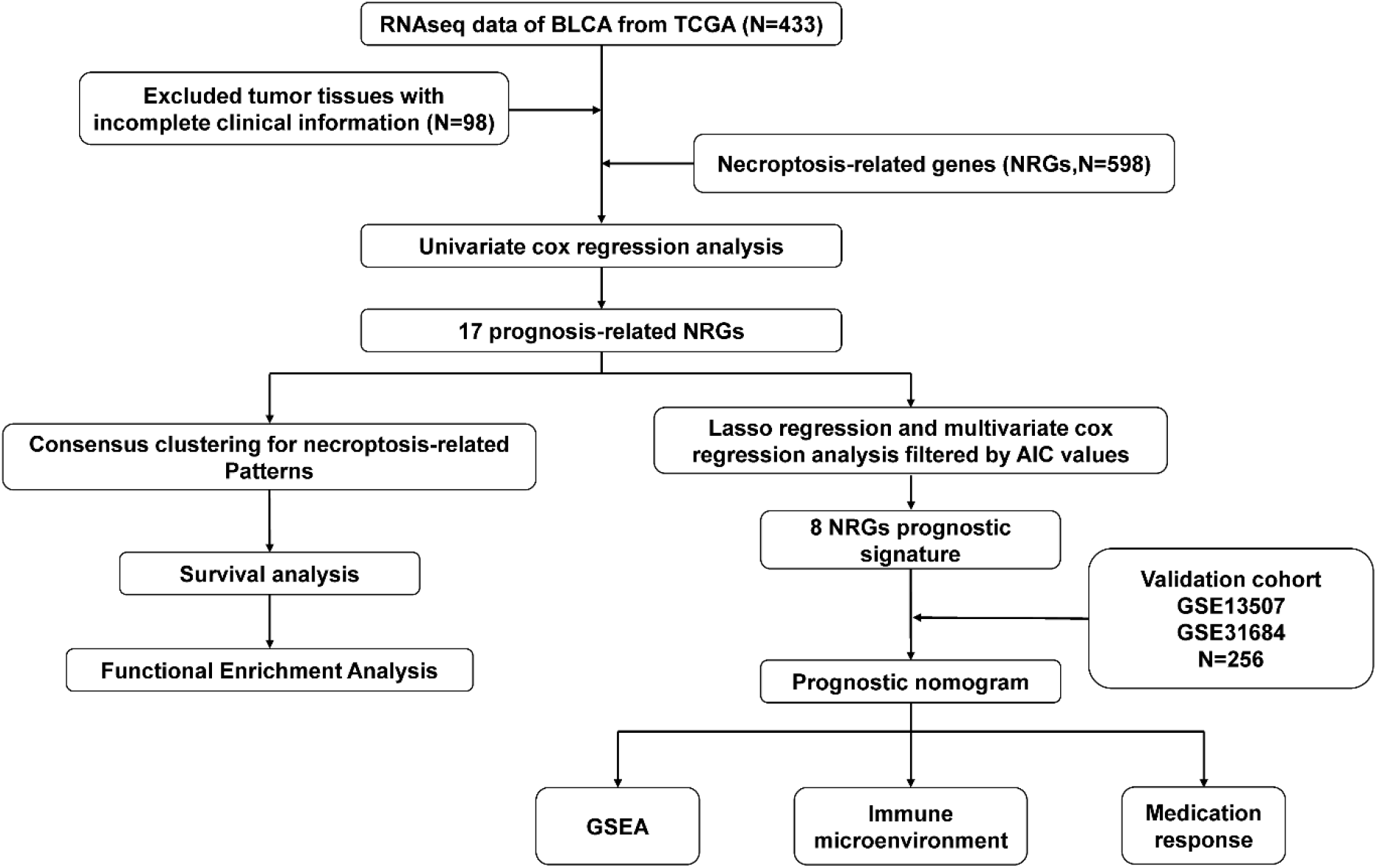
Flow chart of our study.

### 3.1 Consensus clustering of Necroptosis-Related Patterns in BLCA

Through univariate Cox analysis of gene expression and survival data of BLCA patients in the TCGA cohort, we obtained 17 bladder cancer prognosis-related NRGs (**Table 1**). To further explore the correlation between necroptosis-related patterns and survival and clinicopathological data of BLCA patients, we divided BLCA patients into subgroups based on their gene expression. Based on the results of consensus clustering, patients in the TCGA cohort could be divided into two distinct and non-overlapping clusters (**Figure 2A-C**). We performed PCA analysis (**Figure 2D**) on the two clusters, and we found that cluster 1 is mainly distributed in the lower position of the PCA diagram and cluster 2 is mainly distributed in the upper position. The PCA plot showed significant distinction between cluster 1 and cluster 2. Then, the Chi-square test was used to explore whether there were differences in clinicopathological parameters between the two clusters (**Figure 3A**). As shown in the figure, there were significant differences in age (p < 0.01), pathological grade (p < 0.001), pathological stage (p < 0.001), T-stage (p < 0.001), M-stage (p < 0.01), and N-stage (p < 0.05) between the two groups. Besides, advanced clinicopathological parameters are mainly concentrated in cluster 1. When referring to the difference in survival between the two groups, Kaplan–Meier analysis (**Figure 3B**) showed that patients within cluster 1 had worse survival than cluster 2 (p = 0.003). We performed GO and KEGG analyses on DEGs between cluster 1 and cluster 2 to investigate differences in fundamental biological processes between the two clusters. According to the results of the GO analysis, upregulated genes were mainly enriched in humoral immune response, complement activation-classical pathway, extracellular matrix organization, extracellular structure organization, complement activation and external encapsulating structure organization (**Figure 4A, C**). The results of the KEGG analysis showed that these upregulated genes were significantly enriched in staphylococcus aureus infection, phagosome, protein digestion and absorption, TGF-beta signaling pathway, focal adhesion and ECM-receptor interaction (**Figure 4B, D**).

**Table 1.** 17 necroptosis-related genes associated with bladder cancer prognosis identified by univariate analysis.

| Gene | HR | HR.95L | HR.95H | *p* value |
| --- | --- | --- | --- | --- |
| MLKL | 0.660267 | 0.495817 | 0.879261 | 0.004505 |
| UCHL1 | 1.129145 | 1.030412 | 1.237338 | 0.009277 |
| SIRT6 | 0.474079 | 0.319307 | 0.703870 | 0.000214 |
| FASN | 1.347467 | 1.123951 | 1.615432 | 0.001270 |
| GNLY | 0.797385 | 0.692176 | 0.918584 | 0.001711 |
| TRAF5 | 0.640967 | 0.462229 | 0.888820 | 0.007663 |
| FNDC4 | 1.271419 | 1.093672 | 1.478055 | 0.001776 |
| SIRT5 | 0.515172 | 0.334930 | 0.792410 | 0.002535 |
| SRC | 0.766829 | 0.630499 | 0.932636 | 0.007855 |
| ANXA1 | 1.218066 | 1.095015 | 1.354946 | 0.000283 |
| OGT | 0.693606 | 0.540500 | 0.890081 | 0.004040 |
| ID1 | 0.859464 | 0.781176 | 0.945599 | 0.001885 |
| PADI4 | 1.699223 | 1.185152 | 2.436279 | 0.003926 |
| AIM2 | 0.857089 | 0.779962 | 0.941842 | 0.001349 |
| TNFRSF25 | 0.678689 | 0.532755 | 0.864596 | 0.001702 |
| DSTYK | 1.778230 | 1.198776 | 2.637778 | 0.004221 |
| IKBKB | 0.556251 | 0.401806 | 0.770060 | 0.000409 |

**Figure 2.**
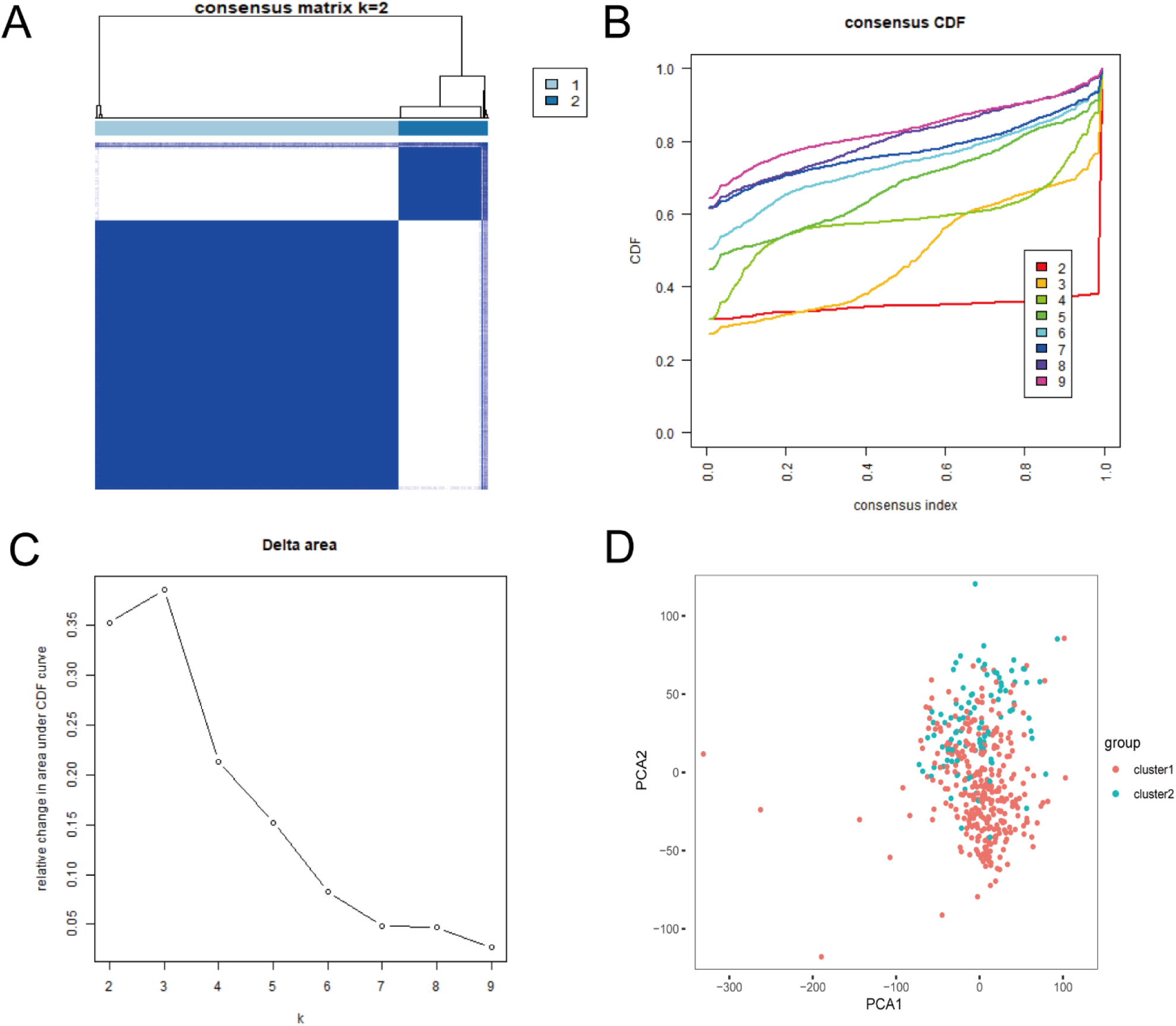
Consensus clustering of BLCA. (A) The correlation between subgroups when consensus matrix number k = 2. (B) Cumulative distribution function (CDF) is displayed for k = 2–9. (C) The relative change in area under the CDF curve for k = 2–9. (D) Principal component analysis (PCA) of the RNA-seq data. Red dots represent cluster 1 and cyan dots represent cluster 2.

**Figure 3.**
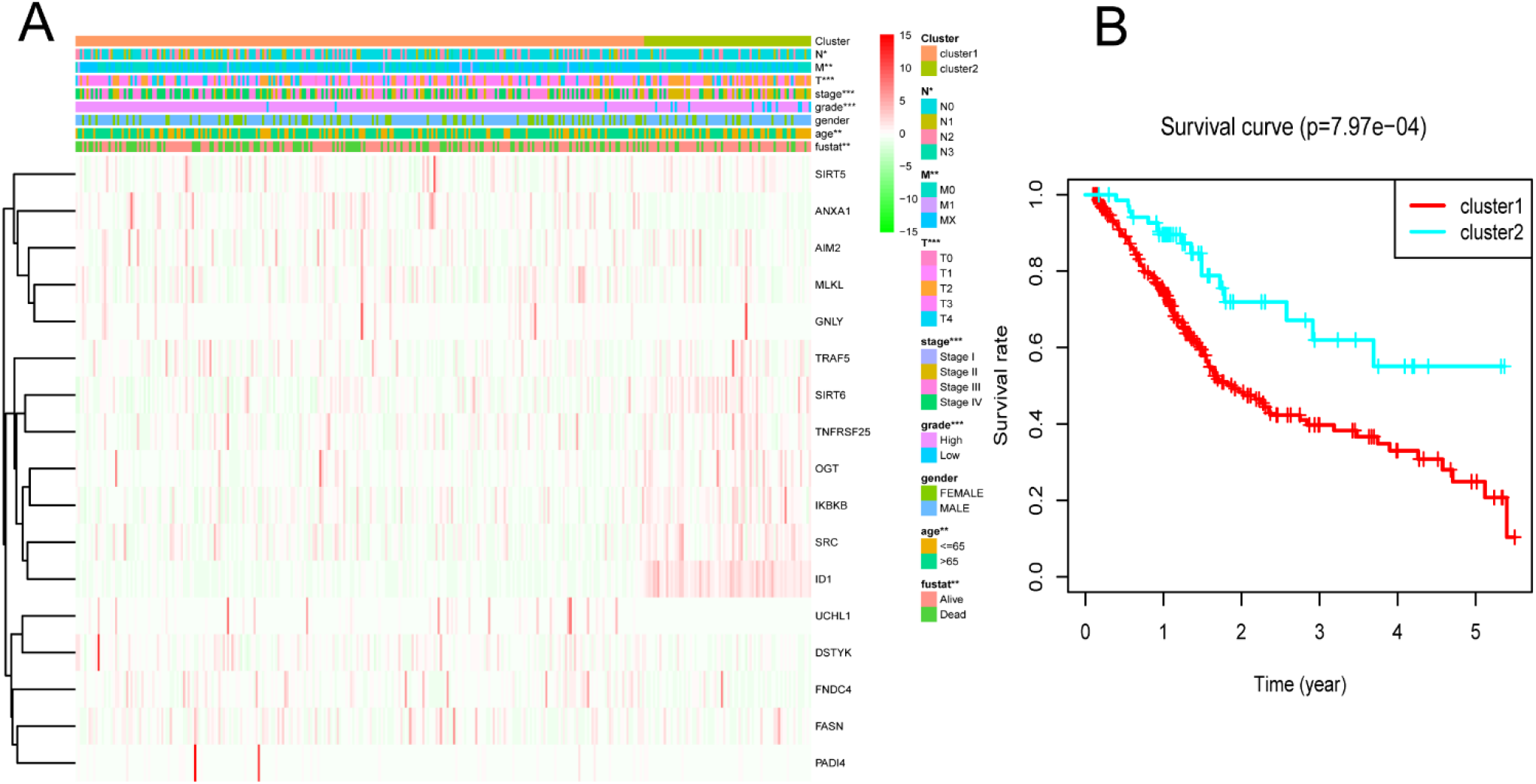
Difference in clinicopathological characteristics and overall survival between cluster 1 and cluster 2. (A) Heatmap and clinicopathological features of these two clusters. (B) Kaplan-Meier survival analysis between cluster 1 and cluster 2. *p < 0.05; **p < 0.01; ***p < 0.001.

**Figure 4.**
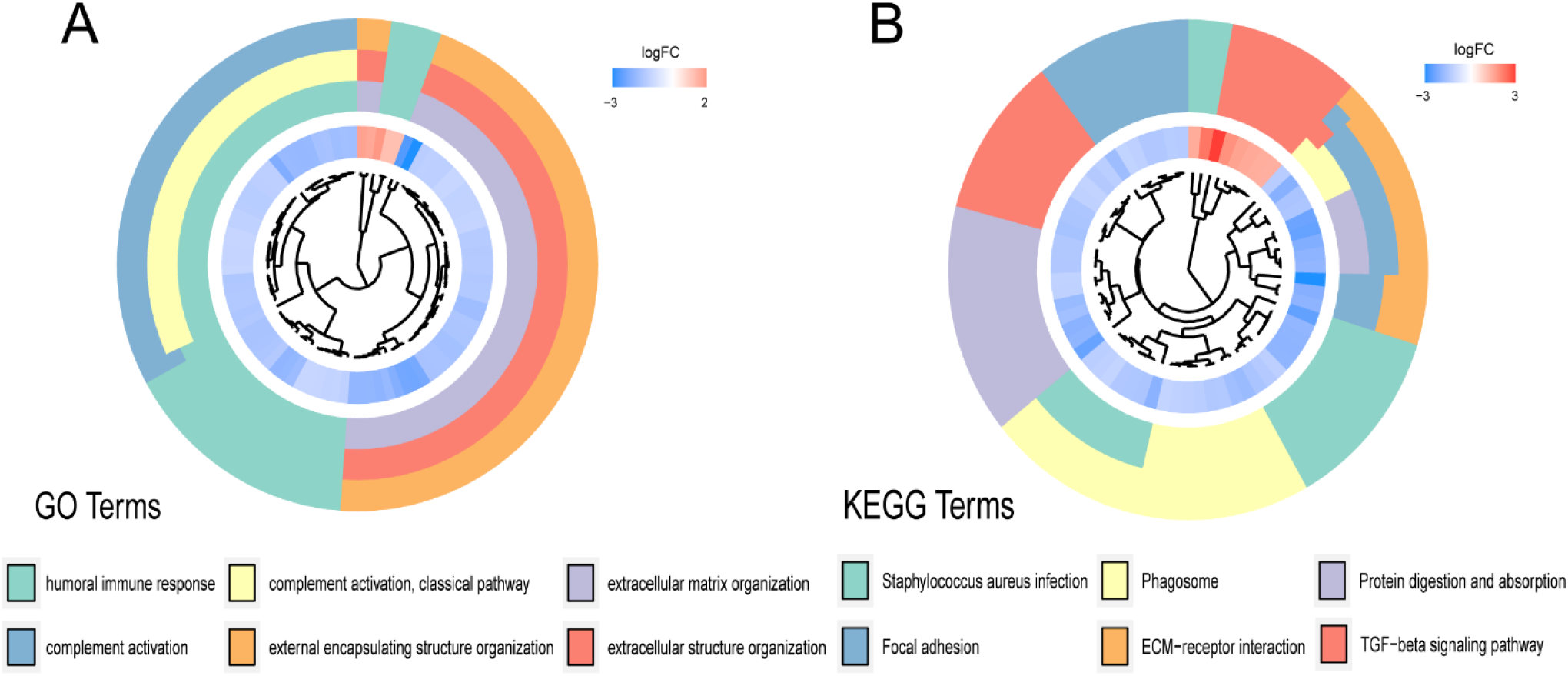
(A, C) Gene ontology (GO) and (B, D) Kyoto Encyclopedia of Genes and Genomes (KEEG) analyses of DEGs between two clusters.

### 3.2 Establishment of the NRGscore in TCGA-BLCA Cohort

To predict the clinical survival rate of BLCA patients accurately and effectively, we establish an NRG prognostic signature from TCGA database. In the previous study, we had obtained 17 BLCA prognosis-related NRGs by univariate Cox regression analysis. We determined NRGs in prognostic signature by LASSO Cox regression (**Figure 5A, B**) analysis and multiple stepwise Cox regression analysis filtered by AIC values. Finally, we identified the optimal prognostic signature containing 8 NRGs, including SIRT6, FASN, GNLY, FNDC4, SRC, ANXA1, AIM2 and IKBKB. We also obtained a quantitative indicator: NRGscore = (−0.1539 × SIRT6 expression) + (0.2289 × FASN expression) + (−0.1683 × GNLY expression) + (0.1683 × FNDC4 expression) + (−0.1851 × SRC expression) + (0.2437 ×ANXA1 expression) + (−0.0918 ×AIM2 expression) + (−0.3567 × IKBKB expression). According to the above calculation formula, we calculate the respective NRGscore for each patient in the TCGA cohort. We divided BLCA patients into low-risk group (N=168) and high-risk group (N=167) based on the median NRGscore to evaluate the prognostic value of NRG signature. Kaplan–Meier analysis showed a worse OS in high-risk group patients relative to low-risk group patients (**Figure 5C**, p < 0.001). Then, we performed the ROC curves to assess the efficacy of the NRG prognostic signature for survival of BLCA patients. As shown in the picture, the area under ROC curves (AUCs) for the 1-, 3-, and 5-year OS were 0.735, 0.768, and 0.745, respectively (**Figure 5D**). **Figure 5E** showed the risk score distribution of patients with BLCA. A dot pot was used to display the survival status of each patient in this cohort (**Figure 5F**). We also evaluated the correlation between NRG prognostic signature and clinicopathological characteristics and show it through a heatmap (**Figure 6**). The high-risk group had advanced pathological grade (p < 0.05), pathological stage (p < 0.01) and N-stage (p < 0.01) than the low-risk group.

**Figure 5.**
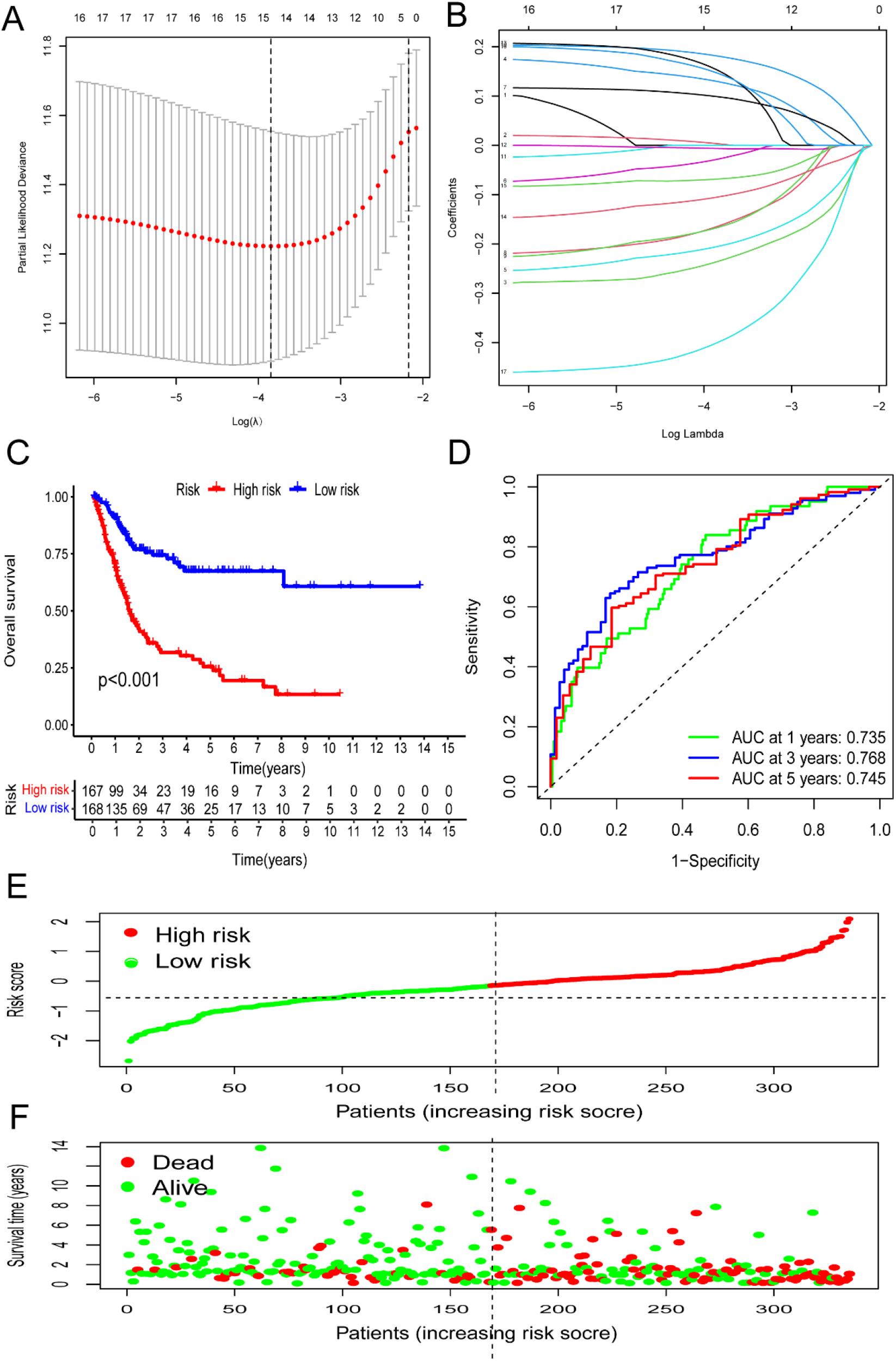
Construction of NRG prognostic signature and prognosis analysis based on the TCGA cohort. (A, B) LASSO Cox regression analysis. (C) Kaplan-Meier survival analysis between NRGscore-defined groups. (D) Time-dependent ROC curves and AUCs at 1-, 3-, and 5-years survival for the0 NRG signature. (E) NRGscore distribution. (F) Survival status map.

**Figure 6.**
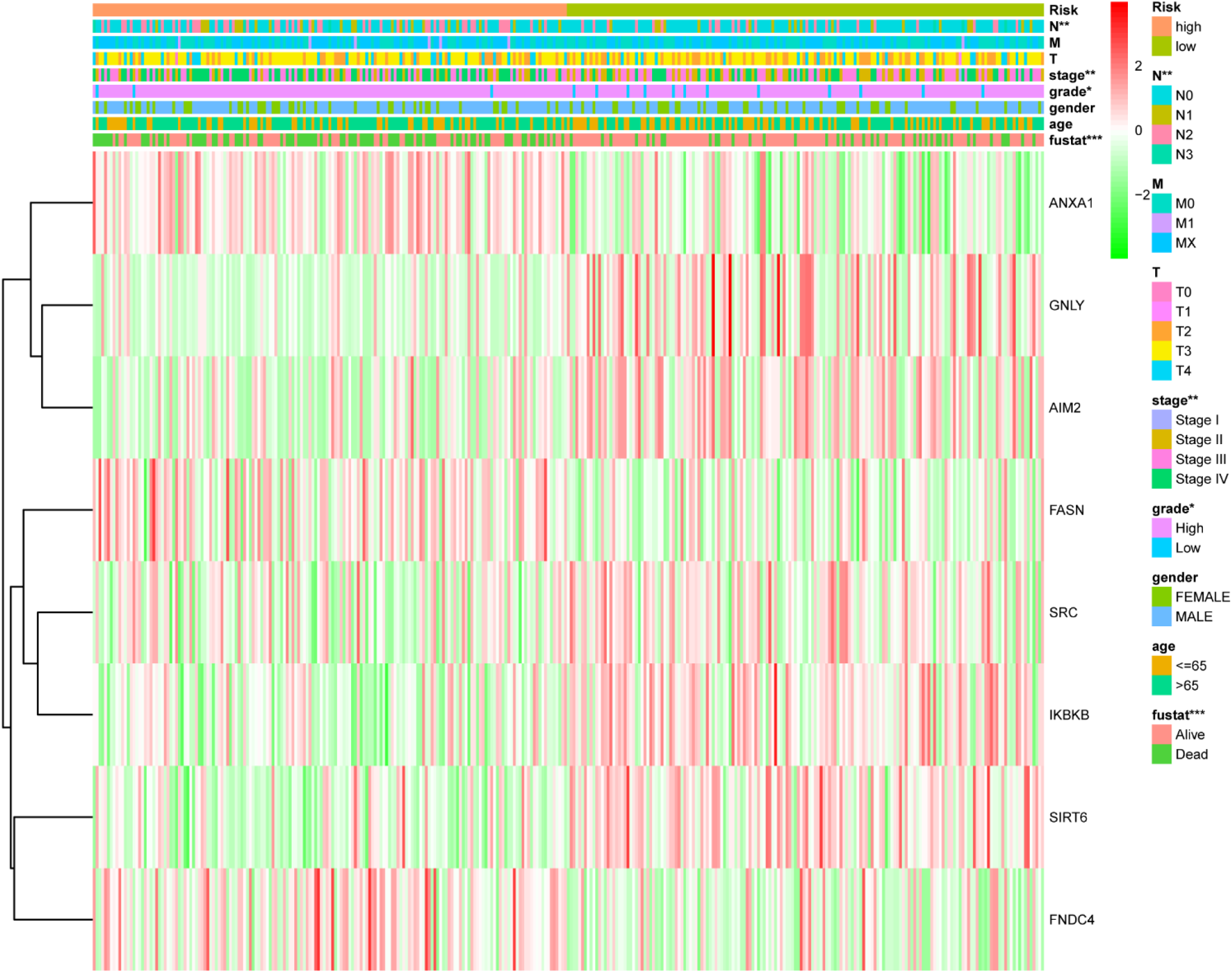
Distribution heat map of NRG signature and clinicopathological variables in the high- and low-risk groups. *p < 0.05, **p < 0.01, ***p < 0.001.

### 3.3 Validation of the Prognostic Signature Using the GEO Database

We used merged GSE13507 and GSE31684 as external validation cohort to estimate the stability and accuracy of the NRG prognostic signature. After excluding two patients who survived less than 30 days, validation cohort included 256 primary bladder cancer patients. BLCA patients were divided into low-risk group (N = 188) and high-risk group (N = 68) according to the cutoff value of the TCGA cohort. We explored the prognostic value of NRG signature in validation and we obtained the results consistent with the TCGA cohort. Patients in the high-risk group had significantly lower OS than those in the low-risk group (**Figure 7A**, p = 0.002). The AUCs for the 1-, 3-, and 5-year OS survival rates were 0.718, 0.606, and 0.577, respectively (**Figure 7B**). The risk score distribution and survival status plots showed similar results to that of the TCGA cohort (**Figures 7C-D)**.

**Figure 7.**
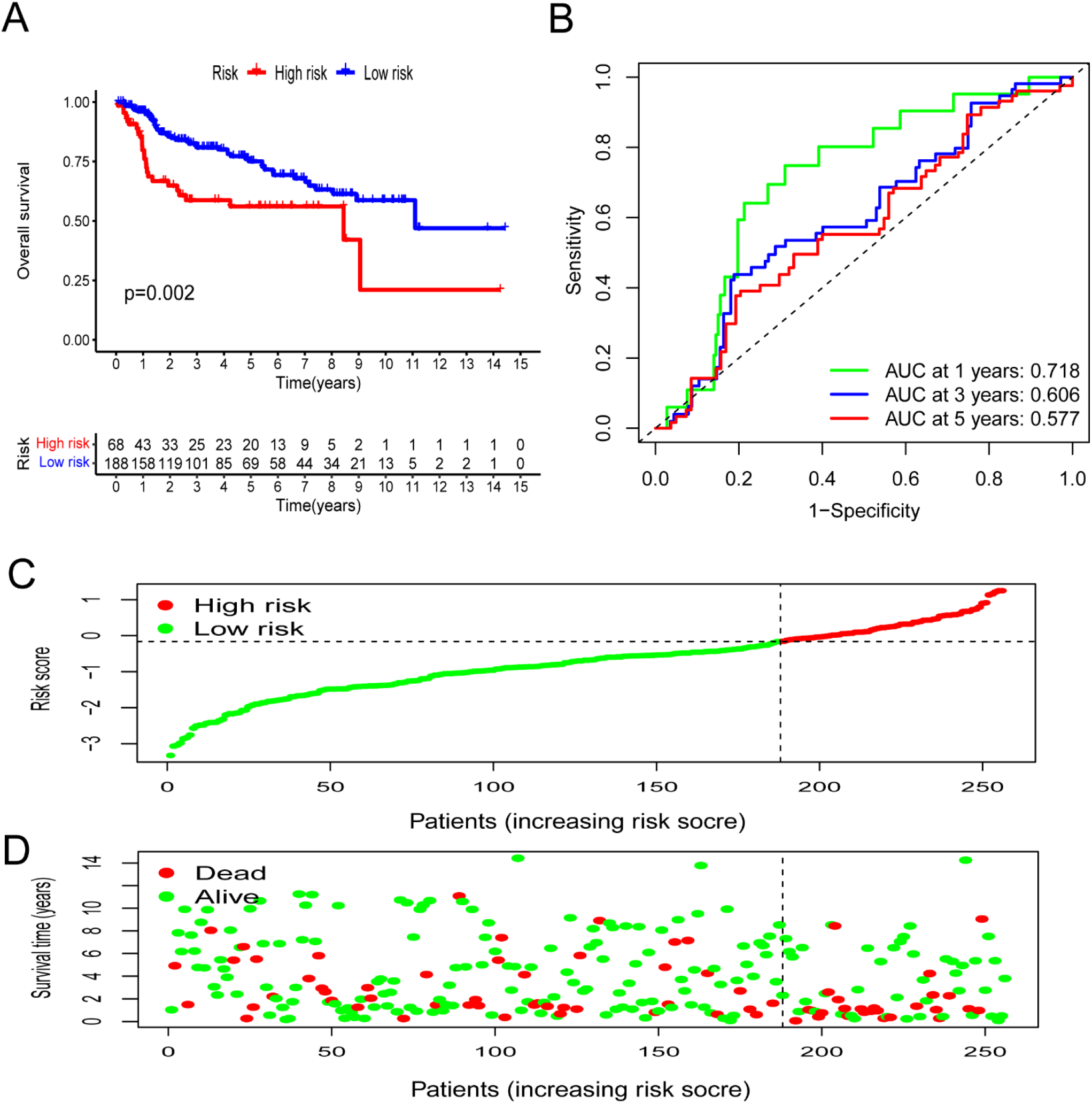
Validation of NRG prognostic signature based on test cohort. (A) Kaplan-Meier survival analysis between NRGscore-defined groups. (B) Time-dependent ROC curves of NRG signature. (C) NRGscore distribution. (D) Survival status map.

### 3.4 Relationship Between the Predictive Signature and the Prognosis of BLCA Patients in Different Clinicopathological Characteristics

We grouped the BLCA patients according to different clinicopathological features to explore the relationship between NRG prognostic signature and each characteristic. Patients were sorted into groups according to age, gender, pathological grade, pathological stage, T-stage, M-stage and N-stage. As shown in **Figure 8**, the OS of BLCA patients in the high-risk group was significantly lower than that in the low-risk group. This suggested that the NRG prognostic signature can accurately predict the prognosis of BLCA patients in different clinicopathological characteristics.

**Figure 8.**
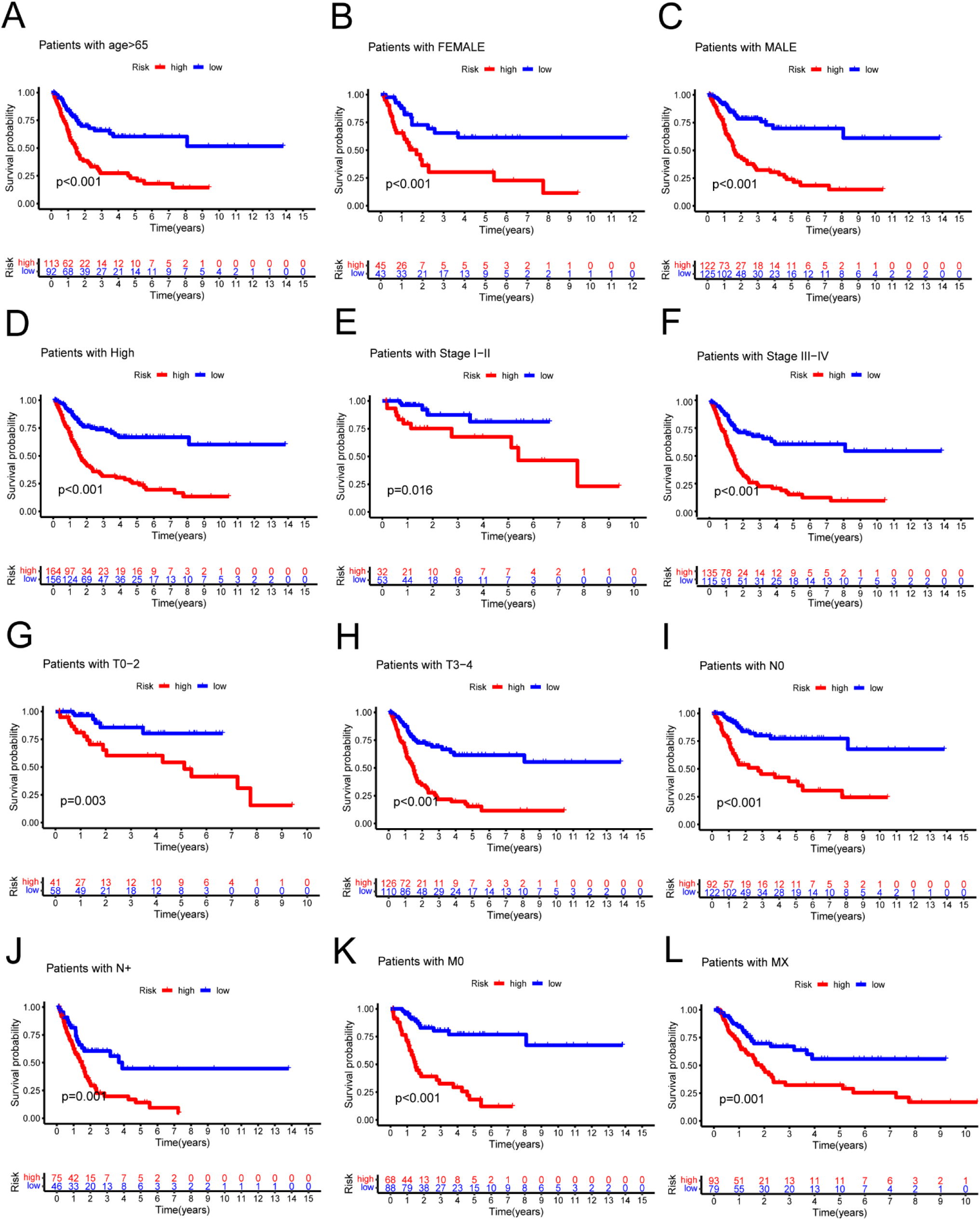
Kaplan-Meier survival curves of high- and low-risk groups among patients sorted according to different clinicopathological features. (A) Age. (B, C) Gender (D) Pathological grade. (E, F) Pathological stage. (G, H) T stage. (I, J) N stage. (K, L) M stage. T, tumor; N, lymph node.

### 3.5 Construction of a Nomogram Model Based on the NRGscore

We performed univariate Cox and multivariate Cox analyses to assess whether NRGscore was an independent prognostic indicator of BLCA patients. Univariate Cox regression analyses (**Figure 9A**) found that age, pathological stage, T-stage, N-stage and NRGscore were prognostic hazard factors. Multivariate Cox regression analyses (**Figure 9B**) pointed out that age (p = 0.024) and NRGscore (p < 0.001) were independent prognostic indicator for BLCA patients. ROC curves of NRGscore and clinicopathological variables was plotted and AUCs results showed that NRGscore was a better independent prognostic factor (**Figure 9C**). Next, we established a novel prognostic nomogram on the basis of age, pathological stage, T-stage and NRGscore (**Figure 10A**) to steadily and accurately predict survival of BLCA patients in TCGA cohort. The calibration curves were used to assess the precision of the nomogram model (**Figures 10B-D**).

**Figure 9.**
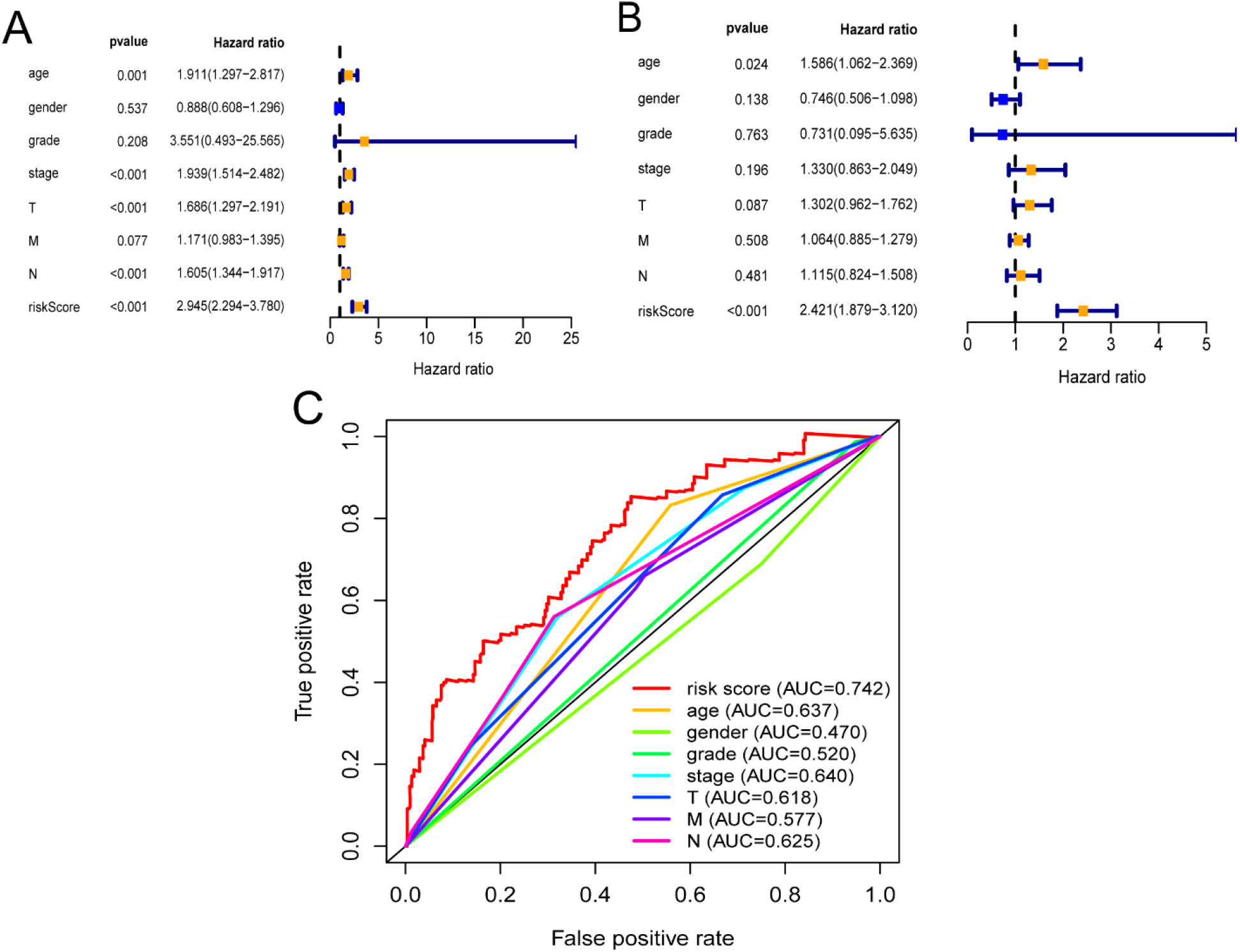
The correlation between NRG signature and the prognosis of BLCA patients. (A) Forest plot for univariate Cox regression analysis. (B) Forest plot for multivariate Cox regression analysis. (C) The ROC curve and AUCs of the risk score and clinicopathological variables. ROC, receiver operating characteristic; AUC, area under the curve; T, tumor; N, lymph node.

**Figure 10.**
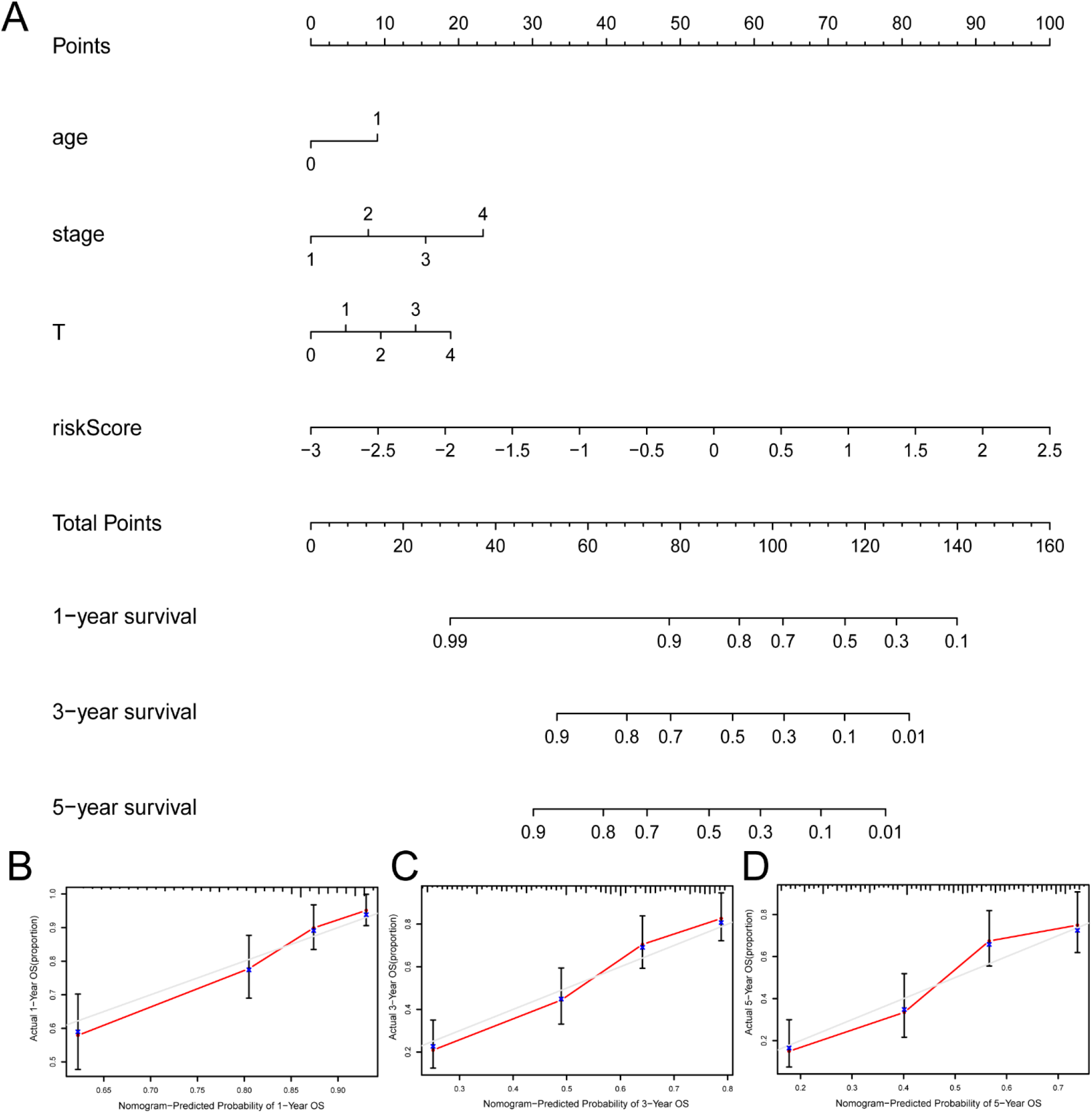
Construction and verification of the nomogram. (A) A nomogram combining clinicopathological variables and risk score predicts 1-, 3-, and 5-years OS of BLCA patients. (B-D) The calibration curves test consistency between the actual OS rates and the predicted survival rates at 1-, 3-and 5-years. N, lymph node; OS, overall survival.

### 3.6 Gene Set Enrichment Analysis

To further comprehend the effect of NRG signature on the biological characteristics of BLCA, we carried enrichment analysis of prognostic signature in GO-GSEA (“pub/gsea/gene_sets/c5.go. v7.5.1.symbols.gmt”) (**Figure 11A, B**). In the high-risk group, we discovered that the enrichment pathway was mainly in various biological regulation, such as organ growth, actomyosin structure organization, cell substrate junction organization, establishment of planar polarity, fibroblast migration. Besides, the enrichment in the localization of nucleus, spindle and the response to platelet-derived growth factor of high-risk group was conspicuous. As for cellular components, high-risk group was enriched in outer membrane. While the low-risk group is mainly concentrated in antigen processing and presentation of endogenous antigen, peptide antigen, and the regulation of innate immune response, signal transduction. Besides, the cellular components of inflammasome complex and U1 snrnp were also enriched in low-risk group. Furthermore, **Figure 11C, D** showed the enrichment of the high-risk and low-risk groups in the Kyoto Encyclopedia of Genes and Genomes (KEGG-GSEA, “pub/gsea/gene_sets/c2.cp.kegg.v7.5.1.symbols.gmt”). In the high-risk group, we found that necroptosis related genes were mainly enriched in the biosynthesis of unsaturated fatty acids and N-glycan. Meanwhile, the enrichment about signaling pathway of TGF-β, WNT and GAP junction, ADHERENS junction was also presented. As for the low-risk group, it was major enriched in multiple diseases such as parkinsons disease, autoimmune thyroid disease, type-1 diabetes mellitus and graft verus host disease. Besides, this group was related to cytosolic DNA sensing pathway, antigen processing and presentation.

**Figure 11.**
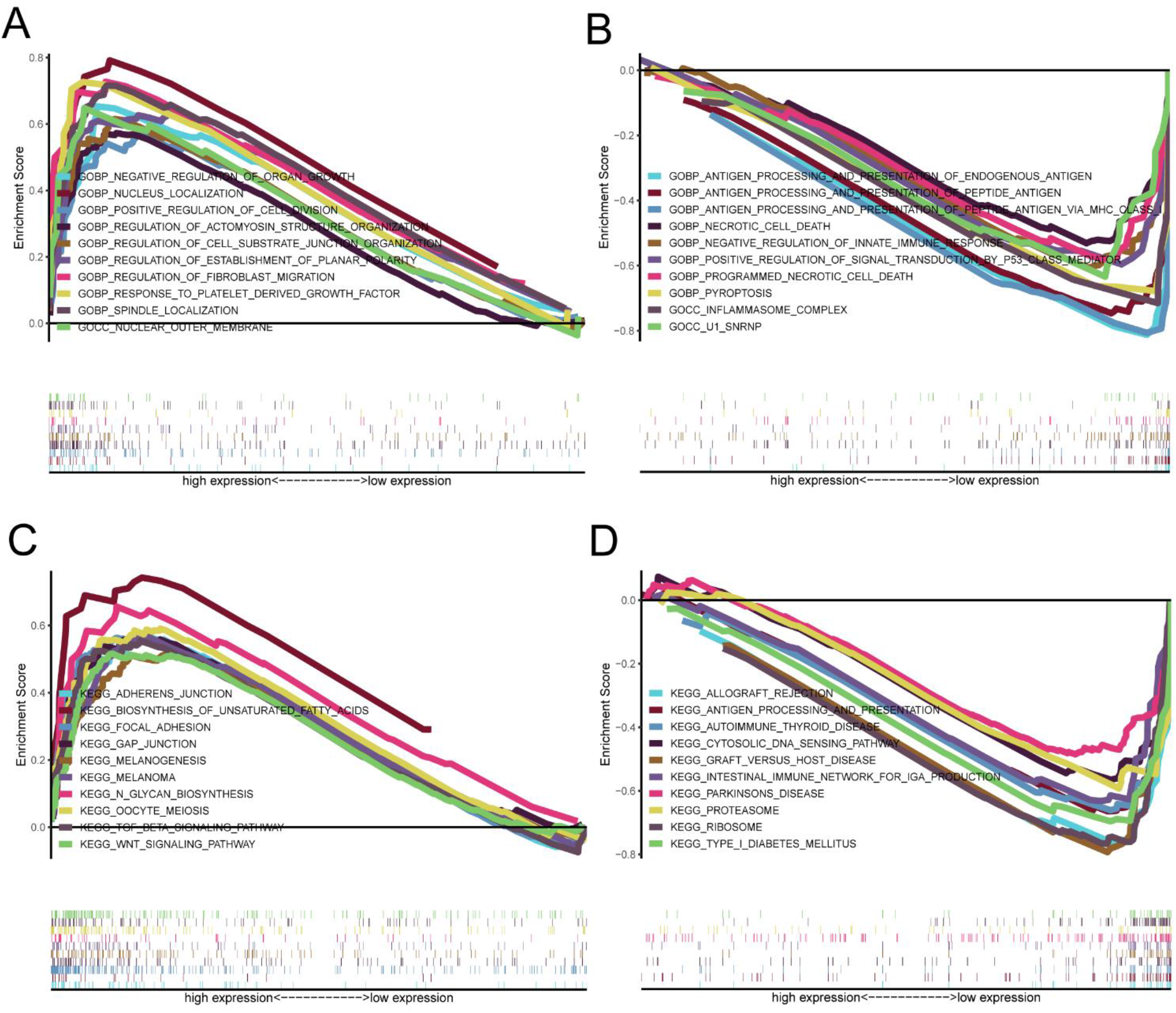
Correlation between NRG signature and biological functions. GO-GSEA results of the high-risk group (A) and low-risk group (B). KEGG-GSEA results of the high-risk group (C) and low-risk group (D).

### 3.7 Correlation Between Tumor Immune Microenvironment and NRGscore

For the importance of checkpoint-based immunotherapy, we made an analysis of expression of immune checkpoints between two-groups in **Figure 12A**. In our research, we discovered that the expression of CD48, CD160, ICOS, CD40, CD40LG, LGALS9, CTLA4, IDO1, TNFRSF14, PDCD1, IDO2, CD27, TNFRSF25, TMIGD2, TIGIT, KIR3DL1, TNFRSF18 and LAG3 in low-risk group was significantly higher than the high-risk group. Meanwhile, the expression of cd200, CD276, NRP1, VTCN1 in low-risk group was less than the high-risk group. In this study, we also put comparative analysis of immune cells to confirm the difference between high-risk and low-risk group (**Figure 12B**). The immune cell score of CD8+T cell, immature dendritic cells (iDCS), NK cells, T helper cells, T helper type 1 (Th1) cells, T helper type 2 (Th2) cells, tumor-infiltrating lymphocyte (TIL) in low-risk group was higher than the high-risk group. We also evaluated the correlation of risk groups with immune-related functions. As shown in **Figure 12C**, the scores of APC_co_inhibition, check-point, cytolytic activity, HLA, inflammation-promoting, MHC-I, parainflammation, T_cell_co_inhibition, T_cell_co-stimulation, and type-1 IFN response in low-risk group were significantly higher than the high-risk group. The score of Type-Ⅱ-IFN_Response in high-risk group was higher than low-risk group. These findings pointed out that NRG prognostic signature had great relationship with tumor immune microenvironment.

**Figure 12.**
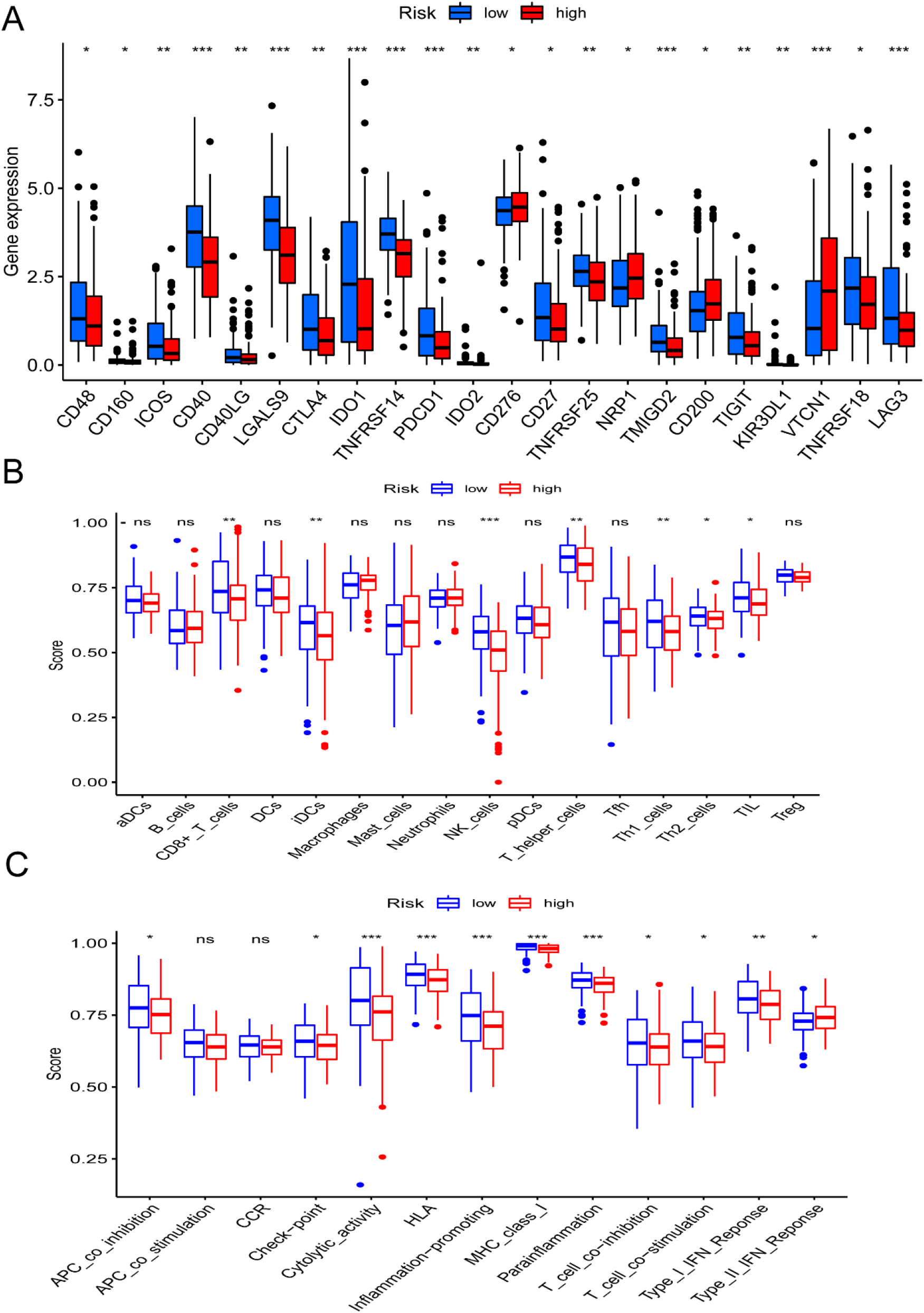
Tumor immune microenvironment of NRG signature. (A) Expression of immune checkpoints among high- and low-risk groups. (B) Infiltration of 22 TIICs in high- and low-risk groups. (C) Immune-related functions in NRGscore-defined groups. *p < 0.05; **p < 0.01; ***p < 0.001; ns = not significant.

### 3.8 Correlation Between the NRG Signature and Drug Sensitivity

The pRRophetic package was used to test the correlation between NRG signature and the sensitivity of different antitumor drugs between high-risk and low-risk groups. We found that the high-risk group had lower IC50s among various small-molecule targeted drugs (**Figure 13A-J**), such as Imatinib, Bexarotene, Midostaurin, FH535, CGP.082996, CMK, PF.4708671, JW.7.52.1, KIN001.135 and Z.LLNle.CHO. These results suggested that patients in the high-risk group may be more sensitive to these drugs, which may be used as a combination therapy drugs for high-risk BLCA patients. We also compared the response to immunotherapy in different risk groups. As shown in **Figure 14A-C**, TIDE score and excusion score were lower in low-risk group, and dysfunction score was lower in high-risk group. When we assessed the immunophenoscore of different risk groups from TICA website, including CTLA4- PD1- group, CTLA4- PD1+ group, CTLA4+ PD1- group and CTLA4+ PD1+ group, we found that low-risk group obtained higher immunophenoscore in all these conditions (**Figure 14D-G**, all p < 0.001). These results suggested that patients in low-risk group may be more sensitive to immunotherapy.

**Figure 13.**
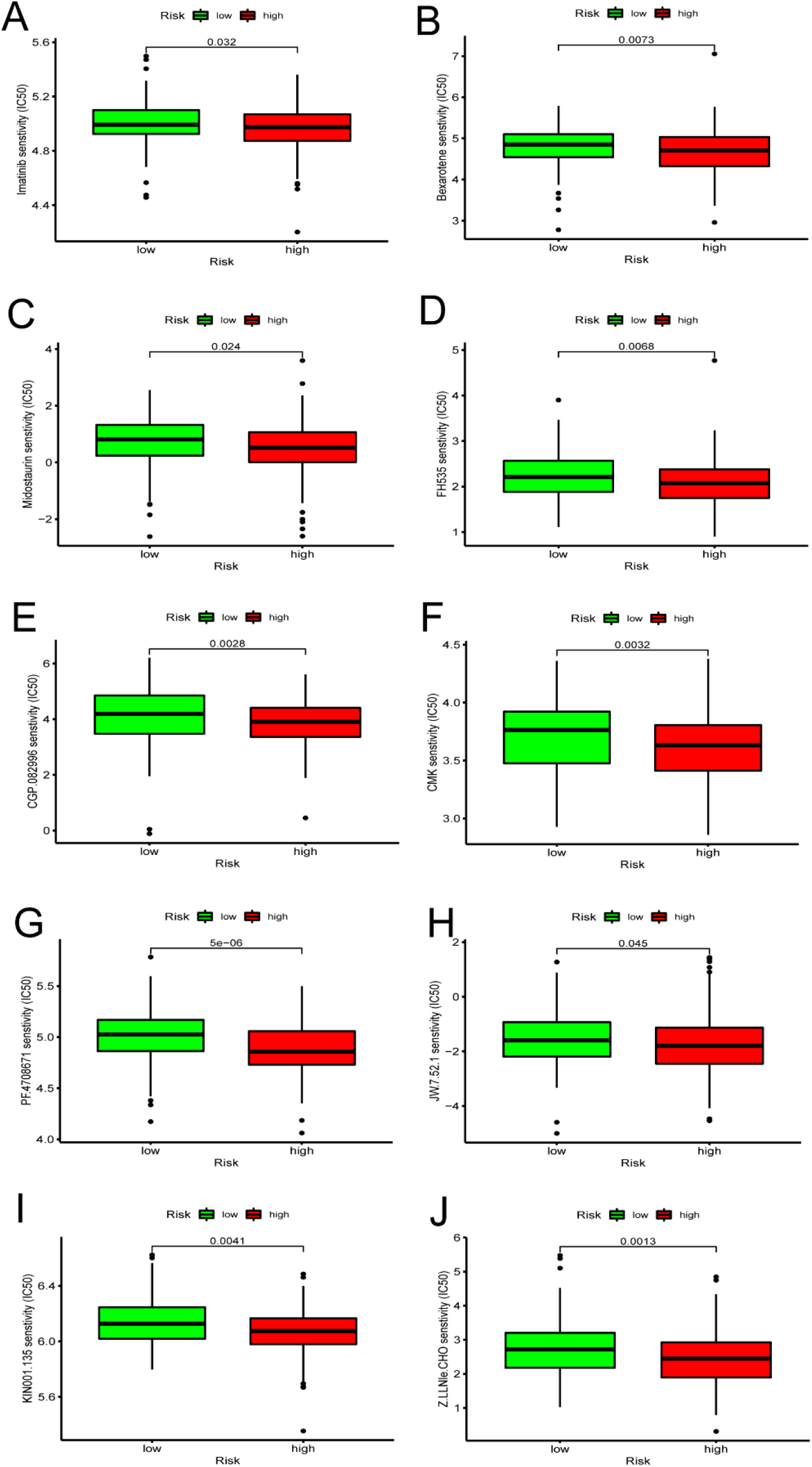
Correlation between the NRG signature and IC50 values of antitumor drugs, including (A) Imatinib, (B) Bexarotene, (C) Midostaurin, (D) FH535, (E) CGP.082996, (F) CMK, (G) PF.4708671, (H) JW.7.52.1, (I) KIN001.135 and (J) Z.LLNle.CHO.

**Figure 14.**
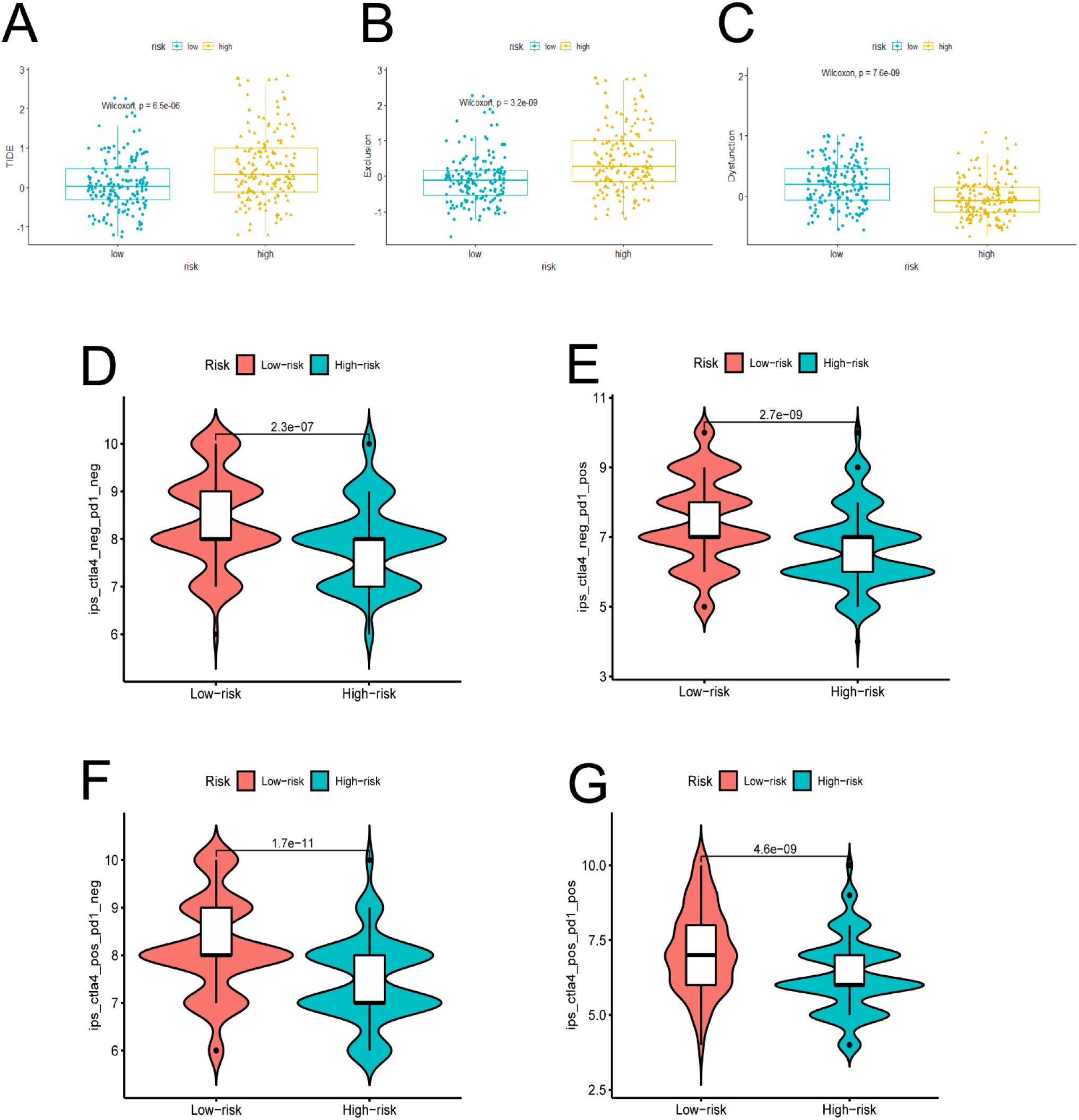
Response of different risk groups to immunotherapy. (A) TIDE score. (B) Exclusion score. (C) Dysfunction score. Immunophenoscore from TICA when (D) CTLA4_neg_PD1_neg, (E) CTLA4_neg_PD1_pos, (F) CTLA4_pos_PD1_neg, (G) CTLA4_pos_PD1_pos. Neg=negative, pos=positive.

## 4 Discussion

Bladder cancer is a highly malignant urological tumor, and even the BLCA patients with lower T stage (T1) have a recurrence rate as high as 23.5% in five years after the treatment of transurethral resection of bladder tumor (TURBT) (18). Although the diagnosis and treatment of BLCA has been improved in recent years, the prognosis of BLCA patients is not satisfactory. Due to the high heterogeneity of BLCA, different BLCA patients often show different prognosis. Previous study suggested that about half of BLCA patients do not benefit from first-line treatment regimens based on combination therapy with gemcitabine and cisplatin, and only 30% BLCA patients achieve remission from ICIs therapy. The tolerance of tumor cells to apoptosis leads to the resistance of cisplatin, leading to the chemotherapy failure (7). Recent studies suggest that promoting immunogenic death may alter tumor microenvironment (TME) and TIL infiltration, and the combination of its inducers and ICIs act synergistically in enhancing antitumor effects (19). Despite accumulating evidence that triggering necroptosis may serve as an alternative method for killing cancer cells, especially those that are not susceptible to apoptosis, research of necroptosis in bladder cancer remains less (20). Therefore, we conducted a comprehensive study of the role of necroptosis in BLCA at present study.

Firstly, we identified 17 NRGs associated with BLCA prognosis by analyzing 598 genes involved in necroptosis pathways using RNA-sequencing results of 335 bladder patients from the TCGA database and constructed two clusters by consensus clustering analysis. Necroptosis C2 showed a significantly better OS survival rate and the proportion of BLCA patients in the early clinicopathological (< T2) stage and low grade of low-risk necroptosis C2 were also significantly higher. More importantly, the differential genes of C2 and C1 were mainly related to the immune response according to GO and KEGG analysis, especially in complement activation and TGF-beta signaling pathway. As we known, complement conducts immune surveillance and inhibits tumor progression through regulation of the immune system and direct killing of tumor cells (21, 22). TGF-beta signaling pathway is a tumor suppressor that inhibits cell proliferation in the early stages of tumors by triggering cell arrest and apoptotic programs in cancer cells (23).

Next, we constructed a novel NRG signature consisting of SIRT6, FASN, GNLY, FNDC4, SRC, ANXA1, AIM2, and IKBKB to predict the survival of TCGA-BLCA cohort (NRGscore). Based on the NRGscore, survival analysis showed that the low-risk group obtained a significantly better OS. Besides, the AUC analysis suggested that NRG has a good predictive value of BLCA patients prognosis. Univariate Cox regression analyses and multivariate Cox regression analyses also confirmed that NRGscore could be considered as an independent prognostic risk, which was significantly proven in two separate external cohorts, including GSE13507 and GSE31684. Among the 8 NRGs included in the NRGscore, ANXA1, FASN, and FNDC4 were up-regulated and GNLY, AIM2, SRC, IKBKB, and SIRT6 were down-regulated in the high-risk group. Recent studies proven that targeting ANXA1 abrogated Treg-mediated immune suppression in triple-negative breast cancer and ANXA1 was considered as a worse prognostic and TME marker in gliomas.(24, 25) FASN is abnormally expressed in lots of tumors and is highly related to tumor migration and invasion (26). FNDC4 promotes the tumor invasiveness of hepatocellular cancer as an extracellular factor to (27). And among these down-regulated genes, Jiang et al. suggested that a signature, including GNLY, was an ideal biomarker for predicting the better prognosis of MIBC patients (28). Fairfax et al. proven that GNLY was also associated with the cytotoxicity and characteristic of effector memory T cells in metastatic melanoma patients (29). Keitaro Fukuda et al. confirmed that AIM2 regulates anti-tumor immunity and is a viable therapeutic target for melanoma (30). Low SIRT6 expression is associated with poorer OS in gastrointestinal cancer (31). However, there are few studies on the relationship between SRC, IKBKB and tumors. In addition, in order to predict the BLCA patient’s OS more accurately and conveniently, we constructed a nomogram combining NRGscore and several clinicopathological features.

Considering that necroptosis is different from apoptosis, which can release cellular contents to trigger a strong immune response, we investigated the correlation between NRGscore and TME. GSEA analysis shown that the low-risk group is mainly concentrated in antigen processing and presentation of endogenous antigen, peptide antigen, and the regulation of innate immune response, signal transduction, which will contribute to the anti-tumor effect (32). In the low-risk group, the infiltration levels of CD8 T cells, NK cells, and iDC cells were significantly increased, which always play important anti-tumor protective roles (33-35). Furthermore, classical immune checkpoints, such as CTLA4, PDCD1, TIGIT, and LAG3 were also enriched in the low-risk group. More importantly, the response to immunotherapy prediction shown that the TIDE score and excusion score were lower in low-risk group, the dysfunction score was lower in high-risk group, which reminds us that the application of immune checkpoint inhibitors in the low-risk group may achieve better therapeutic effects.

We also assessed the value of NRGscore to predict the sensitivity of chemotherapy and targeted agents in BLCA patients. The results shown that imatinib, bexarotene, and midostaurin benefited more in high-risk patients, these drugs are generally used in the treatment of blood cancers. Furthermore, small molecules, including Imatinib, Bexarotene, Midostaurin, FH535, CGP.082996, CMK, PF.4708671, JW.7.52.1, KIN001.135 and Z.LLNle.CHO also had more significant benefits in high-risk patients. Recent research confirmed that targeting necroptosis with small molecules is a promising strategy for cancer therapy, which may be the key to overcoming apoptosis resistance, and has advantages in overcoming apoptosis resistance and stimulating antitumor immunity (36). Therefore, we believe that the NRGscore can help identify better treatment strategies for individual patients with advanced BLCA.

The present study still has some limitations. Firstly, this research is a retrospective study, and the clinical information of BLCA patients inevitably produces biases, and follow-up large, multicenter, prospective studies are needed to further confirm our results. Secondly, the accuracy of the NRGscore model in predicting drug efficacy also needs to be confirmed by clinical trials. Finally, the specific molecular mechanisms of NRG in BLCA that we included in our study remain to be further explored, especially SRC, IKBKB.

## 5 Conclusions

Based on comprehensive analyses, we conducted a NRGscore with excellent performance in assessing the prognosis, clinicopathologic features, TME and therapeutic sensitivity of BLCA patients, which could be utilized as a guide for chemotherapy, ICIs therapy, and combination therapy strategy.

## 7 Author Contributions

ZHZ and NJ developed this idea and designed this research. ZHZ, NJ, and YLZ analysed the data. ZHZ, NJ, THL, YLZ, and TYL wrote the draft of the manuscript. HQG and RY obtained copies of studies and revised the writing. All authors agree to be accountable for the content of the work.

## 8 Acknowledgments

None

## 10 Data Availability Statement

The original datasets for this study can be found in the manuscript/Supplementary material, further information can be obtained by contact the corresponding authors.

